# Six years of clinical herpes simplex virus genotypic acyclovir resistance testing confirms common resistance mechanisms and identifies novel mutations

**DOI:** 10.64898/2026.06.25.734554

**Authors:** Katharine H. D. Crawford, Jared Castor, Katrina LaTurner, Andrew R. Mack, Gregory Pepper, Alexander L. Greninger

## Abstract

Identification of acyclovir-resistant herpes simplex virus (HSV) infections is critical for directing appropriate antiviral therapy, particularly for immunocompromised patients where resistance rates can reach 30%. In 2020, the University of Washington Clinical Virology Laboratory launched the first clinical genotypic HSV drug resistance test in the United States. While genotypic testing offers significantly faster turnaround times than traditional phenotypic assays, interpretation depends on established mutational databases and remains challenging when novel variants are identified. Here, we retrospectively reviewed all HSV acyclovir resistance Sanger sequencing tests performed from January 2020 to November 2025 at this primary national reference laboratory. Mutations identified via clinical sequencing were compared against published databases of HSV UL23 mutations to determine their phenotypic effects. Over the nearly six-year study period, 136 samples were sequenced with a median turnaround time of 10.6 days. Among these, 65 samples (47.8%) harbored acyclovir resistance mutations, including 45 frameshift mutations. Notably, across the 100 samples (73.5%) displaying mutations not known to cause acyclovir resistance at the time of clinical testing, we identified 56 distinct mutations, including 23 without prior characterization. Our national experience demonstrates that genotypic testing accelerates actionable results in clinical practice and confirms that frameshift mutations remain a primary driver of acyclovir resistance. Furthermore, by uncovering these 23 novel variants, this work provides critical targets for future biochemical and phenotypic characterization of HSV UL23 mutations.

**IMPORTANCE:** Acyclovir-resistant herpes simplex virus infections remain an important clinical concern, especially for immunocompromised patients, but testing for acyclovir resistance has traditionally been limited to lengthy and laborious phenotypic testing. Since 2020, the University of Washington Clinical Virology Laboratory has provided genotypic acyclovir resistance testing. This testing provides faster turnaround times but relies on prior mutation characterization to identify resistance making it important to continually evaluate and report identified mutations. The clinical experience with this testing provides valuable insight into resistance mechanisms, identifies common polymorphisms, and highlights novel mutations for further characterization.

## INTRODUCTION

Acyclovir and its prodrug valacyclovir are first-line therapy for herpes simplex virus (HSV) infection treatment or prophylaxis (1, 2). Although uncommon in the immunocompetent population (3, 4), the development of acyclovir resistance remains a significant clinical concern, especially for immunocompromised persons where resistance rates can be up to 30-40%, with the highest rates in hematopoietic stem cell transplant recipients (3, 5). Phenotypic characterization is considered the gold standard for acyclovir drug resistance testing (6), but is limited by availability of HSV culture isolates, assay variability around resistance cut-offs, and lengthy turnaround time (6, 7). Genotypic HSV antiviral resistance testing has been performed for a decade in several European countries and provides an important alternative for clinical testing (6–8).

Compared to phenotypic testing, genotypic drug resistance testing is not reliant on HSV culture, meaning it typically has a faster turnaround time and decreased variability (7, 9). However, result interpretation requires adequate characterization of identified mutations to determine their effect on drug efficacy (7, 10, 11). Acyclovir resistance typically develops from inactivating mutations in the thymidine kinase (*UL23*) gene that is required to phosphorylate acyclovir into its active form (8). Prior to 2020, when the University of Washington (UW) began offering genotypic testing for acyclovir resistance, clinical *UL23* sequencing was primarily performed in Europe and the German Consulting Laboratory for HSV and VZV developed a database of mutation effects to aid in the characterization of HSV UL23 mutations (10, 11). Nonetheless, characterization of UL23 mutations remains incomplete, and clinical sequencing often identifies novel variants that can complicate genotypic drug resistance test interpretation.

To date, the UW Virology Laboratory remains the only clinical laboratory to offer genotypic acyclovir resistance testing in the United States. Here we investigate the first six years of clinical testing to provide updated data on observed mutations and better understand test utilization.

## MATERIALS & METHODS

### Clinical genotypic HSV acyclovir resistance testing

We performed a retrospective review of all HSV drug resistance (HSVDR) testing performed at UW Virology from January 1, 2020 through November 30, 2025. Clinical testing was performed following a clinically validated protocol. HSV *UL23* amplicons were PCR amplified using type-specific primers (**Table 1**) and the following cycling conditions: 98°C for 15 seconds; 40 cycles of: 98°C for 10 seconds, 60°C for 15 seconds, and 68°C for 2 minutes; and a final elongation at 68°C for 7 minutes. Following amplification, samples were run on a 2% agarose gel alongside a 100-base pair (bp) ladder (Thermo Fisher Scientific, California, USA) and visualized using ethidium bromide and a UV Transilluminator to confirm amplification of the expected ∼1,000 bp product. Amplicons were cleaned using ExoSAP-IT (Fisher Scientific, California, USA), and sent to Genewiz (Azenta Life Sciences, New Jersey, USA) for Sanger sequencing (primers available in **Table 1**). Resulting sequences were aligned to the reference *UL23* sequences from HSV-1 strain KOS (KM222721.1) and HSV-2 strain 1998-5562 (MH790662.1) using the latest version of Geneious (currently 2025.2.2) to identify mutations. Characterization of mutations as being associated with drug resistance or not was based on an archived February 2017 version of the Jena database (**Supplemental Data 1 and 2**) (10). All variants not annotated as causing acyclovir resistance in that database were reported as having no evidence of resistance to acyclovir.

**Table 1:**
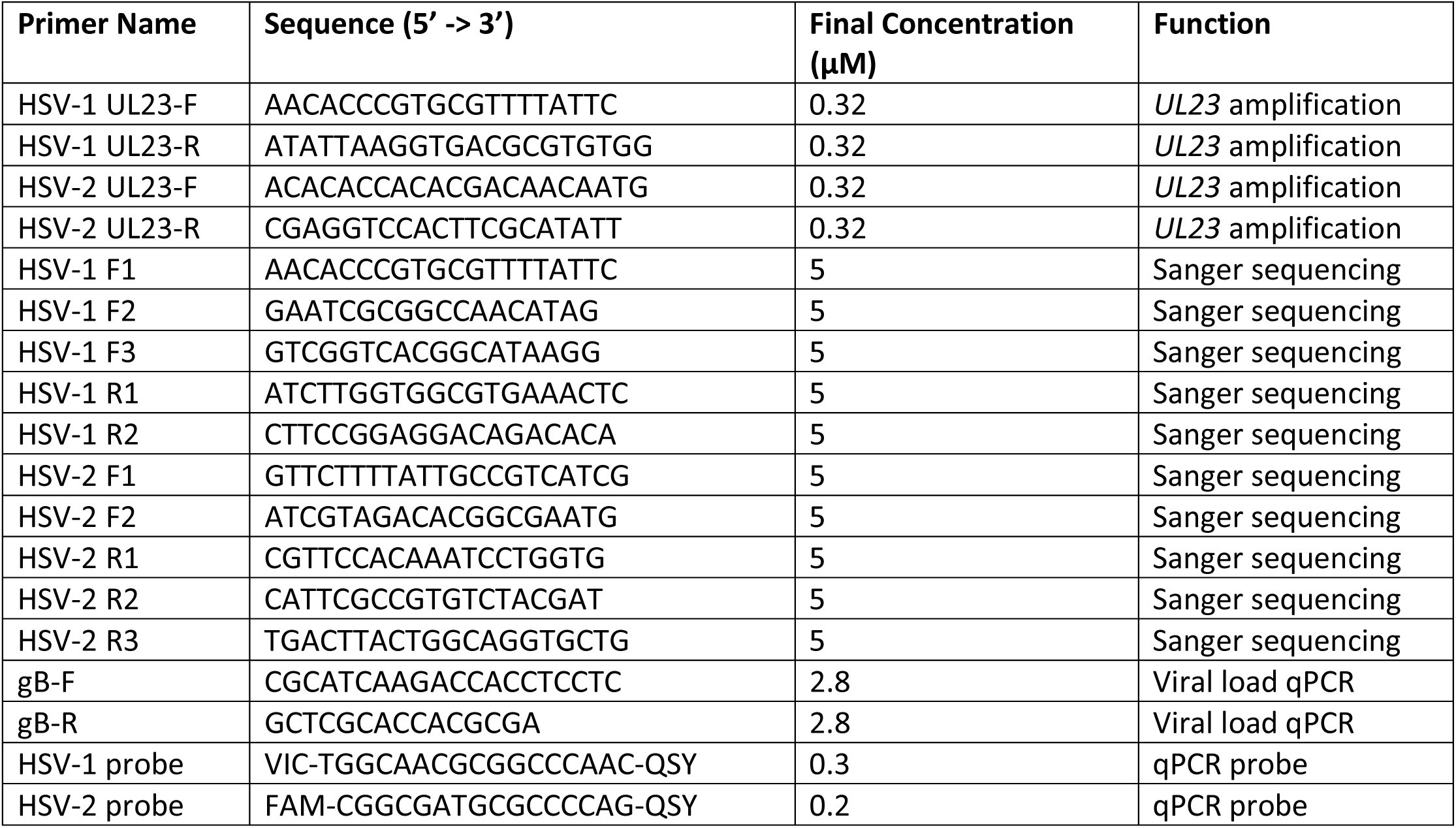
Primer sequences for PCR amplification, sequencing, and viral load testing.

### HSV viral load

To determine HSV viral loads, semiquantitative and quantitative PCR testing was performed according to clinically validated protocols. Semiquantitative testing was performed on swab or cerebrospinal fluid (CSF) specimens using the FDA-cleared Simplexa HSV 1 & 2 Direct assay (Diasorin Inc., Stillwater, MN, USA). Results were reported as detected/not detected with cycle threshold (Ct) values reported for detected samples. Quantitative HSV PCR testing was performed using a clinically validated laboratory developed test (12). Briefly, the HSV *gB* gene was amplified using previously published type-common primers and type-specific probes (**Table 1**) (13). PCR was performed with the QuantiTect Multiplex PCR Mix (Qiagen, Maryland, USA) on an ABI 7500 Real-Time PCR instrument (Thermo Fisher Scientific, California, USA) with the following cycling conditions: 50°C for 2 minutes, 95°C for 15 minutes, followed by 45 cycles of 94°C for 1 minute and 60°C for 1 minute. Ct values were converted to copies/mL using a standard curve of HSV-1 *gB* or HSV-2 *gB* plasmids (Blue Heron, Eurofins Genomics, Washington, USA) at known copy number (1.0×10^8^ -1.0×10^4^ with four 1:10 serial dilutions). Standard curve samples were calibrated with the Acrometrix HSV-1 and HSV-2 plasma panels (Thermo Fisher Scientific, California, USA) to ensure accurate quantification.

### Sequence Analysis

All UL23 protein sequences for HSV-1 and HSV-2 were downloaded from GenBank on January 31, 2026, including 1,572 sequences for HSV-1 and 681 for HSV-2. Sequences were filtered by length to include HSV-1 sequences with 330-377 amino acids and HSV-2 sequences with 306-377 amino acids. This ensured retention of samples with common truncations due to poor annotation of the initial start codon. Filtered sequences were aligned using mafft (v7.525). After an initial alignment, one additional HSV-1 UL23 sequence (HSV1_7 strain, QCX34919.1) was removed due to substantial non-homology through the second half of the sequence and the remaining samples were realigned. The final HSV-1 alignment contained 1,478 sequences and the final HSV-2 alignment contained 661 sequences. The frequency of amino acid mutations at sites where clinical mutations were identified was determined after excluding gaps or unknown (‘X’) residues at each site.

## RESULTS

From January 2020 through November 2025, we sequenced 136 samples for HSV acyclovir resistance testing from 120 persons. There were 89 (65.4%) HSV-1 sequences and 47 (34.6%) HSV-2 sequences (**Table 2**). The median age of patients with sequenced samples was 47 (range: 0-77 years) with a near-equal distribution of male and female patients (female: 61, male: 58, unknown: 1). Only 23 (19.1%) patients were from the UW Medicine system, meaning additional demographic information was unavailable for most patients. Of the 23 UW patients tested, 15 were hematopoietic stem cell transplant recipients (including one person living with HIV) and 11/15 (73.3%) of these patients had at least one resistant sample; 3 patients had a hematologic malignancy without a stem cell transplant, none of whom had acyclovir resistance; 3 were persons living with HIV, two of whom had at least one resistant sample; one patient had a history of solid organ transplant and had acyclovir resistance; and one had an isolated ocular infection without acyclovir resistance.

**Table 2:** Number of samples for each HSV type with no mutations, known resistance mutations, other mutations, or both types of mutations compared to the reference sequences (KM222721.1 [HSV-1] and MH790662.1 [HSV-2]).

| HSV Type | No Mutations | Resistance Mutations Only | Other Mutations Only (incl. 3 ATP-binding site muts) | Both Resistance & Other Mutations | Total |
| --- | --- | --- | --- | --- | --- |
| HSV-1 | 11 | 15 | 37 | 26 | 89 |
| HSV-2 | 4 | 6 | 19 | 18 | 47 |
| **Either** | **15** | **21** | **56** | **44** | **136** |

Of the 136 sequenced samples, 65 (47.8%) had known resistance mutations identified, while 15 (11.0%) had no mutations from the HSV-1 or HSV-2 reference sequences (**Table 2**). Three HSV-1 samples had mutations (P57F + H58Y, G61E, and G61R) detected in the ATP-binding site where other mutations had previously been associated with acyclovir resistance (10, 11). These samples were reported with an interpretation of “No Evidence of Resistance,” but the mutations were reported as potential mutations of concern in the “drug resistance mutations” result field. All three samples also had additional mutations not known to be associated with drug resistance.

The most common drug resistance mutations were frameshifts with 45 of the 65 resistant samples (69.2%) having frameshift mutations leading to early translation termination (**Table 3**, **Figure 1**). Frameshift mutations most commonly occurred at sites of homopolymer runs of guanines (fs146/fs147 in HSV-1/HSV-2) or cytosines (fs185/fs186 in HSV-1/HSV-2) (14, 15). Three HSV-1 sequences had two resistance-associated substitutions identified (W88R + D162N, T245M + A189V, and E83K + A93V). Of note, one nonsense mutation (Q8stop) identified in an HSV-2 sequence was reported as a resistance mutation, though this mutation is not associated with resistance in HSV-1 (11, 16). In-frame stop codon mutations within the first 33 amino acids of HSV-1 UL23 are not expected to cause acyclovir resistance as *in vitro* thymidine kinase activity is unaffected by truncation of the first 33 amino acids of HSV-1 UL23, likely due to the presence of a potential secondary translation initiation site at site 46 (16, 17). However, HSV-2 does not have a possible secondary translation initiation site until site 70, after the ATP-binding site. Therefore, we cannot extrapolate the effects of early in-frame stop mutations from in HSV-1 to HSV-2 and further investigation is required to better understand UL23 truncations in HSV-2.

**Figure 1.**
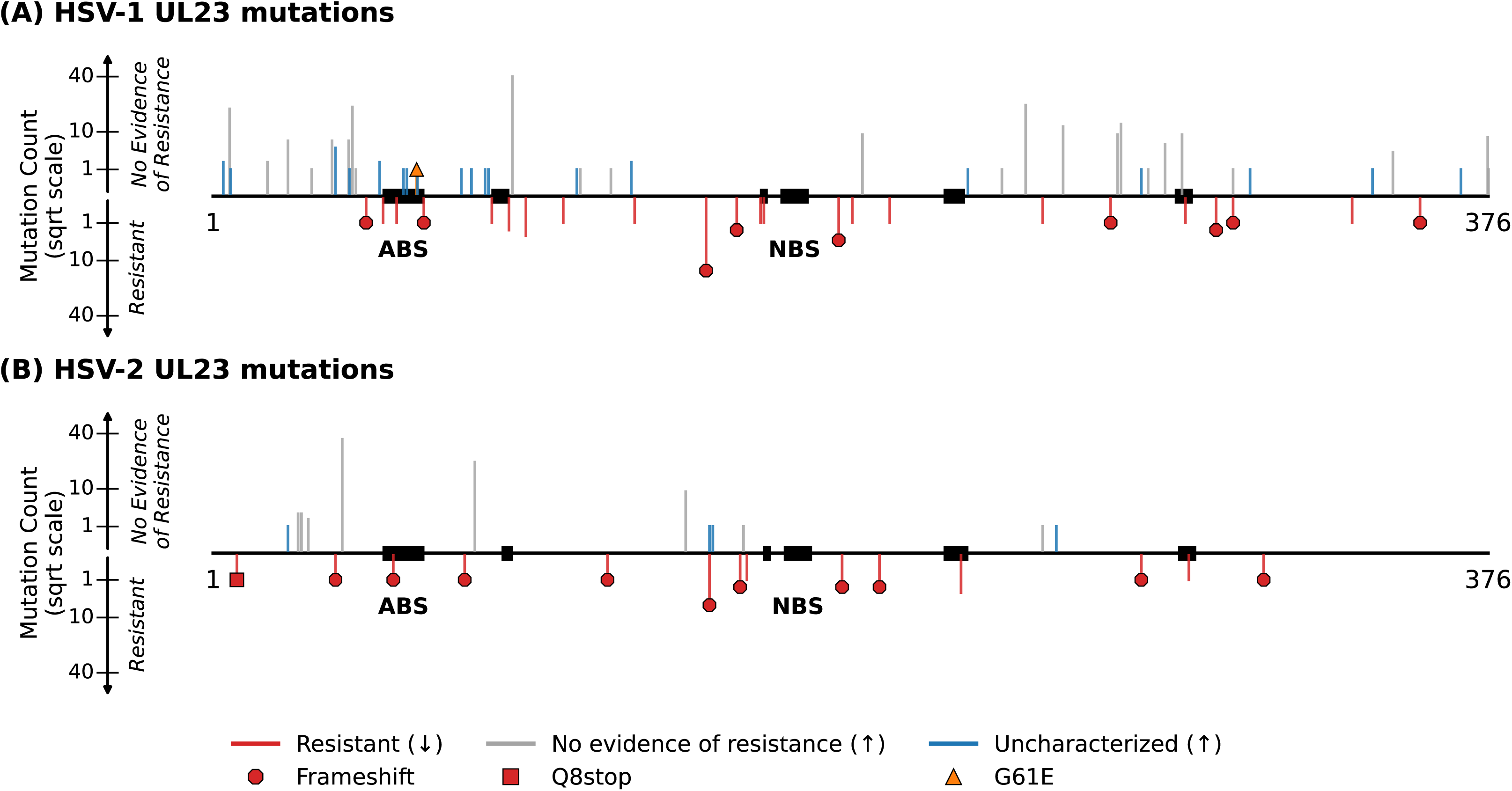
Location and frequency of mutations observed in HSV-1 UL23 (A) and HSV-2 UL23 (B). Mutations are oriented and colored by whether they have been characterized previously and association with acyclovir resistance. Line length corresponds to the number of times that mutation was observed. If multiple mutations were identified at one site, the bars are slightly offset but overlapping. The Q8stop and G61E mutations are highlighted. Conserved regions of UL23 are represented by black boxes. ABS, ATP-binding site. NBS, nucleotide-binding site.

**Table 3:**
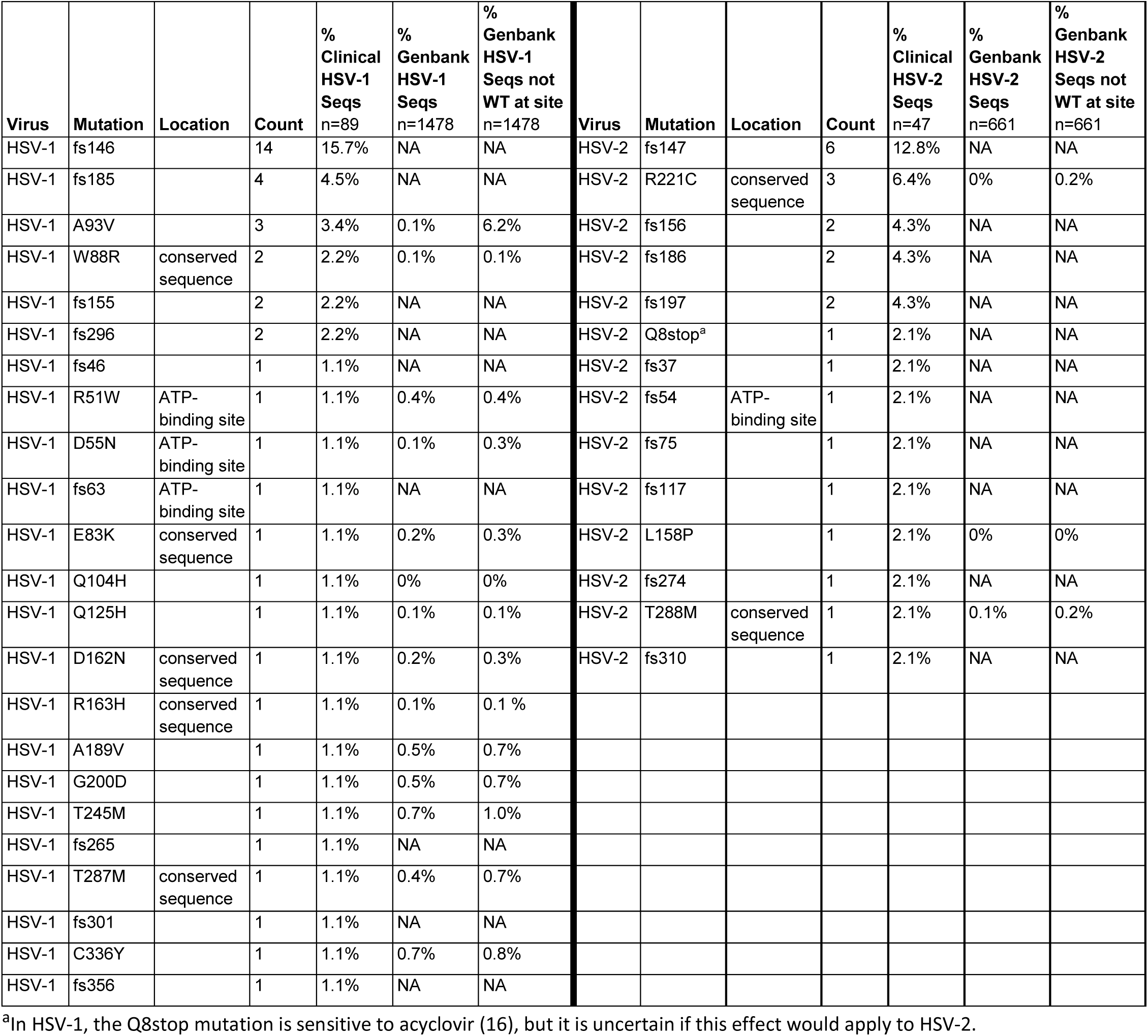
All identified known acyclovir resistance-associated mutations in descending order of frequency for HSV-1 and HSV-2. Location categorization was based on (10). All missense mutations were previously identified as resistance-associated mutations (10, 11). WT = wildtype.

| Virus | Mutation | Location | Count | % Clinical HSV-1 Seqs n=89 | % Genbank HSV-1 Seqs n=1478 | % Genbank HSV-1 Seqs not WT at site n=1478 | Virus | Mutation | Location | Count | % Clinical HSV-2 Seqs n=47 | % Genbank HSV-2 Seqs n=661 | % Genbank HSV-2 Seqs not WT at site n=661 |
| --- | --- | --- | --- | --- | --- | --- | --- | --- | --- | --- | --- | --- | --- |
| HSV-1 | fs146 |  | 14 | 15.7% | NA | NA | HSV-2 | fs147 |  | 6 | 12.8% | NA | NA |
| HSV-1 | fs185 |  | 4 | 4.5% | NA | NA | HSV-2 | R221C | conserved sequence | 3 | 6.4% | 0% | 0.2% |
| HSV-1 | A93V |  | 3 | 3.4% | 0.1% | 6.2% | HSV-2 | fs156 |  | 2 | 4.3% | NA | NA |
| HSV-1 | W88R | conserved sequence | 2 | 2.2% | 0.1% | 0.1% | HSV-2 | fs186 |  | 2 | 4.3% | NA | NA |
| HSV-1 | fs155 |  | 2 | 2.2% | NA | NA | HSV-2 | fs197 |  | 2 | 4.3% | NA | NA |
| HSV-1 | fs296 |  | 2 | 2.2% | NA | NA | HSV-2 | Q8stop<sup>a</sup> |  | 1 | 2.1% | NA | NA |
| HSV-1 | fs46 |  | 1 | 1.1% | NA | NA | HSV-2 | fs37 |  | 1 | 2.1% | NA | NA |
| HSV-1 | R51W | ATP-binding site | 1 | 1.1% | 0.4% | 0.4% | HSV-2 | fs54 | ATP-binding site | 1 | 2.1% | NA | NA |
| HSV-1 | D55N | ATP-binding site | 1 | 1.1% | 0.1% | 0.3% | HSV-2 | fs75 |  | 1 | 2.1% | NA | NA |
| HSV-1 | fs63 | ATP-binding site | 1 | 1.1% | NA | NA | HSV-2 | fs117 |  | 1 | 2.1% | NA | NA |
| HSV-1 | E83K | conserved sequence | 1 | 1.1% | 0.2% | 0.3% | HSV-2 | L158P |  | 1 | 2.1% | 0% | 0% |
| HSV-1 | Q104H |  | 1 | 1.1% | 0% | 0% | HSV-2 | fs274 |  | 1 | 2.1% | NA | NA |
| HSV-1 | Q125H |  | 1 | 1.1% | 0.1% | 0.1% | HSV-2 | T288M | conserved sequence | 1 | 2.1% | 0.1% | 0.2% |
| HSV-1 | D162N | conserved sequence | 1 | 1.1% | 0.2% | 0.3% | HSV-2 | fs310 |  | 1 | 2.1% | NA | NA |
| HSV-1 | R163H | conserved sequence | 1 | 1.1% | 0.1% | 0.1 % |  |  |  |  |  |  |  |
| HSV-1 | A189V |  | 1 | 1.1% | 0.5% | 0.7% |  |  |  |  |  |  |  |
| HSV-1 | G200D |  | 1 | 1.1% | 0.5% | 0.7% |  |  |  |  |  |  |  |
| HSV-1 | T245M |  | 1 | 1.1% | 0.7% | 1.0% |  |  |  |  |  |  |  |
| HSV-1 | fs265 |  | 1 | 1.1% | NA | NA |  |  |  |  |  |  |  |
| HSV-1 | T287M | conserved sequence | 1 | 1.1% | 0.4% | 0.7% |  |  |  |  |  |  |  |
| HSV-1 | fs301 |  | 1 | 1.1% | NA | NA |  |  |  |  |  |  |  |
| HSV-1 | C336Y |  | 1 | 1.1% | 0.7% | 0.8% |  |  |  |  |  |  |  |
| HSV-1 | fs356 |  | 1 | 1.1% | NA | NA |  |  |  |  |  |  |  |

<sup>a</sup>In HSV-1, the Q8stop mutation is sensitive to acyclovir (16), but it is uncertain if this effect would apply to HSV-2.

All amino acid substitutions known to cause HSV drug resistance had frequencies <1% in GenBank UL23 sequences (**Table 3**). The low prevalence of HSV resistance mutations suggests they arise primarily under selective pressure from acyclovir therapy and that many different possible mutations can disrupt UL23 activity. The HSV-1 mutation A93V was the only drug-resistance mutation identified at a site where >1% of sequences contained non-wildtype amino acids. A93V has previously been reported as having uncertain significance (3, 18), but was characterized as a resistance mutation following isolation from a patient with acyclovir resistance without other known resistance mutations (11, 19). We required all substitutions we reported as causing acyclovir resistance to be previously characterized or to cause early translation termination either through a frameshift or an in-frame stop codon at any point in HSV-2 UL23 or after the first 33 amino acids of HSV-1 UL23.

For the 100 samples that had mutations not known to be associated with acyclovir resistance, each sequence frequently had multiple mutations with an average of 3.1 mutations per sequence (range: 1-10). We identified 56 distinct HSV UL23 mutations from these samples (44 in HSV-1 and 12 in HSV-2). Thirty-three mutations (25 in HSV-1 and 8 in HSV-2) (58.9%) had been previously reported in the HSV UL23 mutation databases (10, 11). All previously reported mutations were associated with sensitivity to acyclovir except for HSV-1 G61E, which was reported as resistant to acyclovir in the most recently updated database (11). G61E was already reported as a potential drug resistance mutation, but this new classification allows us to formally report sequences with that mutation as resistant.

We identified 23 mutations – 19 in HSV-1 UL23 and 4 in HSV-2 UL23 – in our samples that were not previously characterized. We examined the location of these mutations and their frequency in publicly available sequences in GenBank to provide additional context for these uncharacterized mutations (**Table 4**, **Figure 1, Figure S1**) (20). Three of these previously uncharacterized mutations (P57F, H58Y, and G61R) were in the ATP-binding site, making them more likely to be resistance mutations (15). Interestingly, the H58Y mutation is present at a frequency of 6.2% among GenBank sequences, which is substantially higher than any known resistance mutation and potentially indicates that mutation is more likely to be a polymorphism despite its location in the ATP-binding site. One other mutation (Y4H) had a frequency >1% in GenBank (4.3%), also suggesting it is likely a polymorphic mutation. Examining mutation frequency in available databases or looking at the location of these mutations on the structure of UL23 (**Figure 1, Figure S1**) can help generate hypotheses about mutation effects, but further biochemical or phenotypic studies are needed to fully characterize all 23 uncharacterized mutations.

**Table 4:**
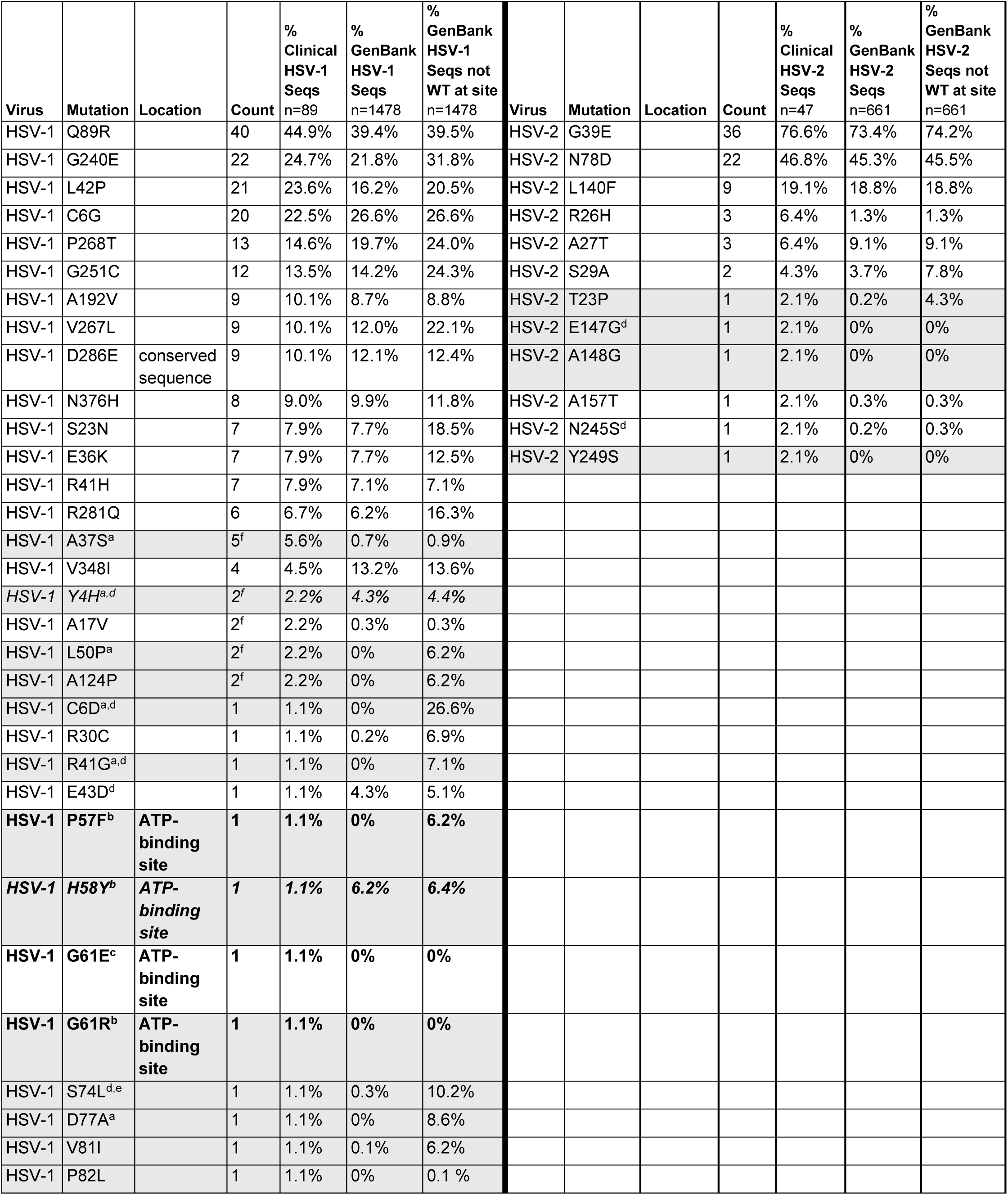
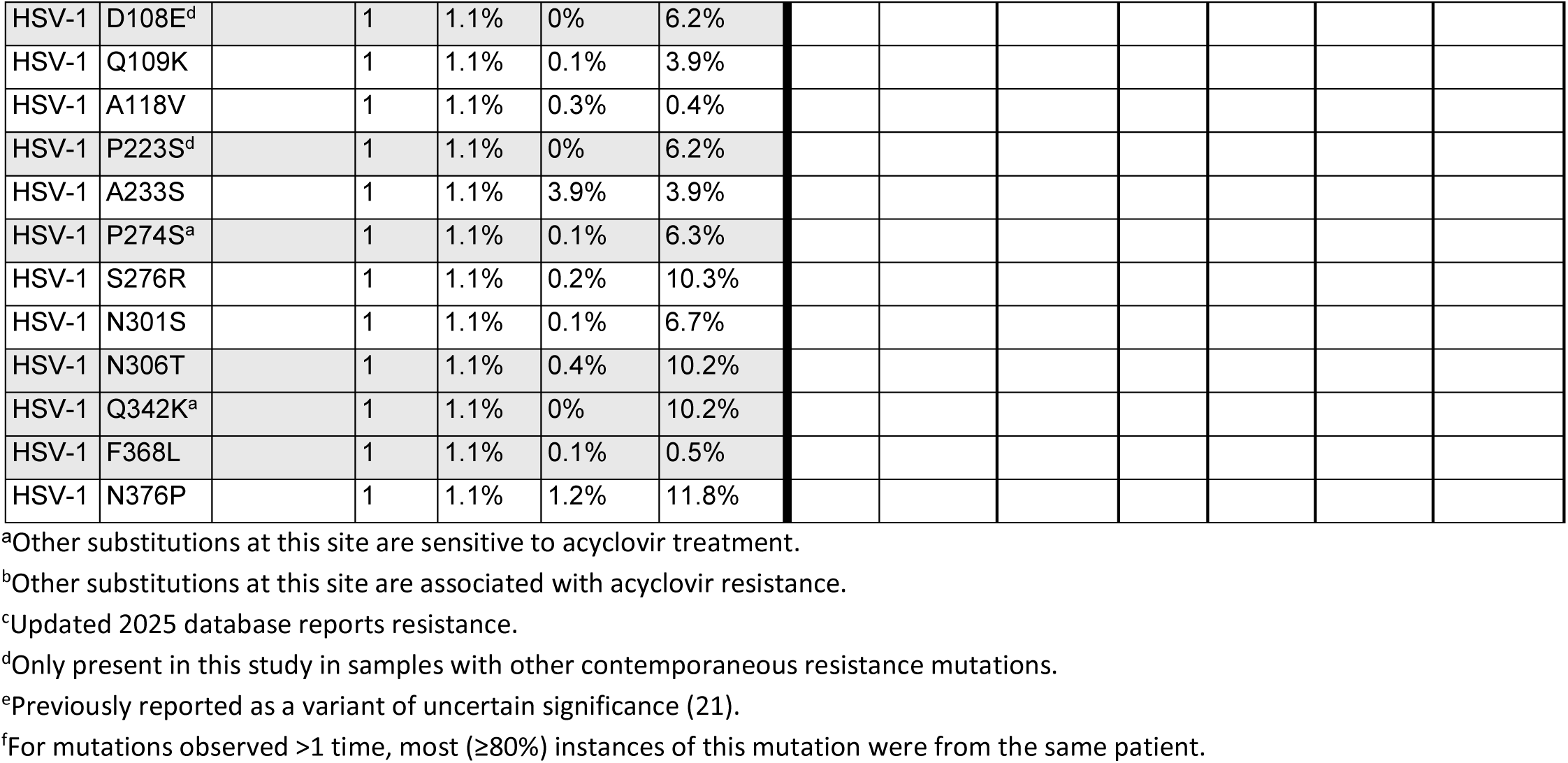
All identified mutations not known to be associated with acyclovir resistance at the time of clinical testing in descending order of frequency for HSV-1 and HSV-2. One sequence could have multiple mutations. Bolded rows identify mutations in the ATP-binding site that were clinically reported as potential resistance mutations. Grey-shaded rows identify mutations that were not previously reported in the HSV UL23 mutation databases (10, 11). Italicized rows were mutations not previously characterized, but present at a frequency >1% in GenBank. Location was determined based on (10) and, if not listed, was non-conserved sequence.

### Patients with multiple sequences

There were 12 patients with multiple sequenced samples with an average of 2.3 sequences per person (max 4) collected over an average of 5 months (range: 1 day - 2.3 years) (**Table 5**). Eight people (66.7%) had no difference in sequence interpretation across their samples, including one person who had an additional mutation not associated with acyclovir resistance identified on subsequent sequencing. Three people (25.0%) had resistance develop over time. Two of these people were UW patients with prior hematopoietic stem cell transplantation and the third sample came from an outside client meaning clinical history was unavailable. One person (8.3%) had an acyclovir resistance frameshift mutation identified in their first sample, but not in a second sample collected two weeks later. This patient had recently received a hematopoietic stem cell transplant and was being actively treated with foscarnet following the initial acyclovir-resistant sample.

**Table 5:** Sequence information from persons with multiple sequences. Patients marked with a * had resistance develop across their samples and the patient marked with ** lost resistance. When sample source was vague (e.g., “swab”), it is not known whether all samples from one person came from the same site or not. BAL = bronchoalveolar lavage.

| Patient | Type | Number of Seqs | Specimen Sources (in order) | Timespan (months) | Initial Result | Initial Drug Resistance Mutations | Initial Other Mutations | Final Result | Final Drug Resistance Mutations | Final Other Mutations |
| --- | --- | --- | --- | --- | --- | --- | --- | --- | --- | --- |
| A* | HSV-1 | 2 | Plasma, Mouth swab | 9.5 | No Evidence of Resistance | Not Detected | G240E | Resistant | Q104H | G240E |
| B | HSV-2 | 3 | Anal skin tag, Swab (x2) | 27.6 | No Evidence of Resistance | Not Detected | G39E N78D | No Evidence of Resistance | Not Detected | G39E N78D |
| C* | HSV-1 | 3 | Swab (x3) | 1.2 | No Evidence of Resistance | Not Detected | G240E | Resistant | fs146 | Not Detected |
| D | HSV-1 | 2 | Swab, Plasma | 1.7 | Resistant | fs296 | Y4H, G240E | Resistant | fs296 | Y4H, G240E |
| E | HSV-2 | 2 | Finger (x2) | 3.5 | Resistant | fs197 | G39E | Resistant | fs197 | G39E, E147G |
| F* | HSV-1 | 4 | Swab (x4) | 9.9 | No Evidence of Resistance | Not Detected | A37S, L42P, Q89R, N301S, F368L | Resistant | fs146 | A37S, L42P, Q89R |
| G | HSV-1 | 2 | Swab (x2) | 1.6 | No Evidence of Resistance | Not Detected | A17V, A124P | No Evidence of Resistance | Not Detected | A17V, A124P |
| H | HSV-1 | 2 | BAL, Swab | 0.2 | No Evidence of Resistance | Not Detected | C6G, L42P, L50P, Q89R | No Evidence of Resistance | Not Detected | C6G, L42P, L50P, Q89R |
| I | HSV-1 | 2 | Swab (x2) | 3.0 | Resistant | fs146 | Q89R | Resistant | fs146 | Q89R |
| J | HSV-1 | 2 | Swab (x2) | 0.6 | No Evidence of Resistance | Not Detected | Q89R, G251C, P268T | No Evidence of Resistance | Not Detected | Q89R, G251C, P268T |
| K** | HSV-2 | 2 | Swab (x2) | 0.5 | Resistant | fs147 |  | No Evidence of Resistance | Not Detected | Not Detected |
| L | HSV-1 | 2 | Eye swab (x2) | 0.03 | No Evidence of Resistance | Not Detected |  | No Evidence of Resistance | Not Detected | Not Detected |

### Operational considerations

Although we sequenced 136 samples, we were surprised to receive many orders for samples that had insufficient HSV viral loads for sequencing (**Table 6**). From January 1, 2020 through November 30, 2025, 138 (45.2%) of the 305 unique orders for HSV drug resistance testing had too low of an HSV viral load either to produce an amplicon for *UL23* sequencing (Low Viral Load) or to obtain valid sequence results after attempting sequencing (No Sequence Obtained) (**Table 6**). Thirty-one (10.2%) orders were canceled for operational reasons (e.g., incorrect order, incorrect sample type, etc.) and were not considered further in our analyses (**Table 6**).

**Table 6:** Result tally and turnaround times for all HSV drug resistance orders from January 1, 2020 through November 30, 2025.

| Result Category | Count | Turnaround Time in Days<br>median (min-max) |
| --- | --- | --- |
| Low Viral Load | 86 | 6.2 (0.8-22.7) |
| No Sequence Obtained | 52 | 7.9 (2.2-19.7) |
| No Evidence of Resistance | 71 | 10.4 (3.0-37.9) |
| Resistant | 65 | 10.7 (3.2-52.8<sup>a</sup>) |
| Other Cancel Code (e.g., duplicate order,<br>canceled by provider, incorrect order, etc.) | 31 | 4.9 (0.8-25.0) |

<sup>a</sup>The sample with a 52.8-day TAT was submitted prior to going live with clinical testing and held until testing was available. TAT from test availability was 11 days.

Samples that were unable to be sequenced came from 124 different persons, including 14 who also had a successfully sequenced sample. Patients with samples that were unable to be sequenced had similar demographics as patients with successfully sequenced samples—median age 48 years, 47.5% female, 50.8% male, and 1.6% unknown sex. Clinical testing was performed once a week with turnaround times of 10.6 days if amplification was successful and 7.1 days if no valid amplicon was produced (**Table 6**).

Despite specifying a minimum copy number (1,000 copies/mL) for testing and requesting any available HSV viral load information when we received samples for clinical testing, only 71 of 274 samples (25.9%) had quantitative (30 samples) or semiquantitative (41 samples) HSV viral load information available to us prior to HSV drug resistance testing. We ran additional quantitative or semiquantitative HSV qPCR on 158 samples, yielding viral load data for 229 samples. Surprisingly, 60 (26.2%) of these samples had no detectable HSV. Samples with undetectable HSV had a similar distribution of internal UW (21.6%) and outside (78.3%) orders as successfully sequenced samples (23.5% internal, 76.5% outside).

Consistent with our previously determined quantitative threshold for successful HSV drug resistance testing of 1,000 copies/mL, no samples that were successfully sequenced had <1,000 copies/mL (**Figure 2A**). Similarly, no samples with successful sequencing had Ct values in the semiquantitative assay ≥35 (**Figure 2B**). For samples with both quantitative and semiquantitative data, results were well-correlated. A viral load of 1,000 copies/mL corresponded to a Ct of 34, and a Ct of <35 was determined to be needed to proceed with sequencing (**Figure 2C**). Of the 150 samples with known HSV viral loads and ≥1,000 copies/mL or Ct <35, 124 (82.7%) were successfully sequenced. Samples that were not successfully sequenced were most often close to the threshold or had insufficient sample volume for sequencing after performing ordered viral load testing.

**Figure 2.**
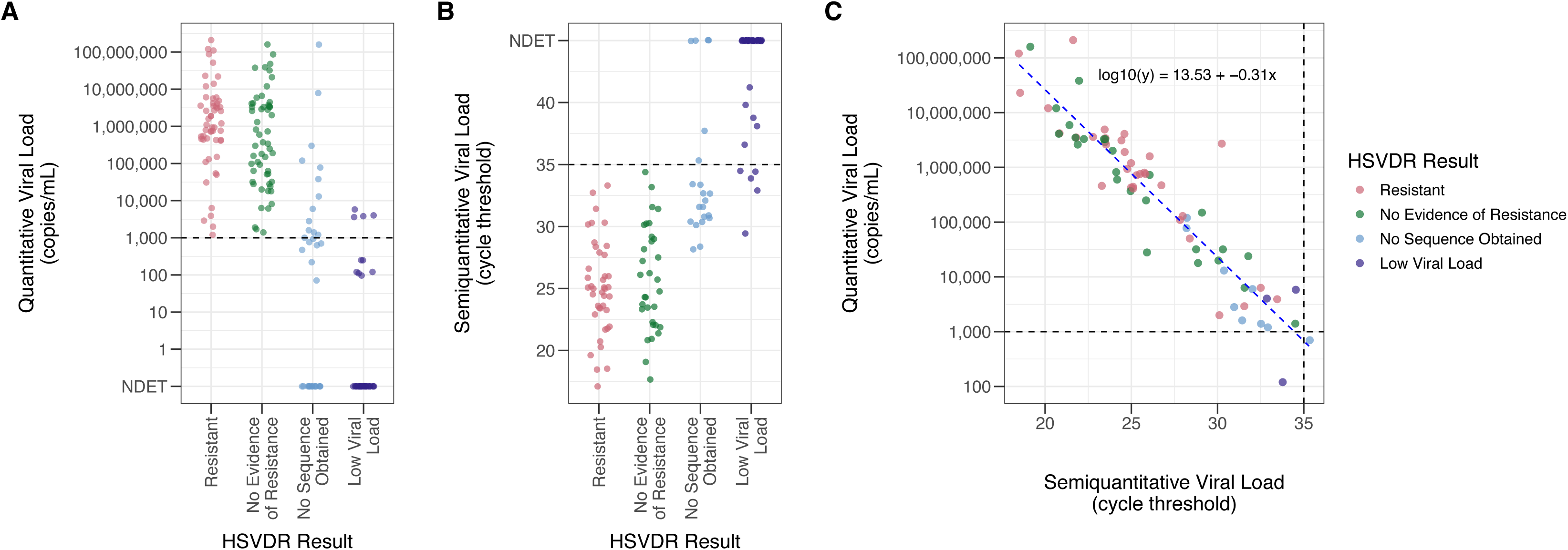
HSV viral loads by result for samples submitted for sequencing. A) All results with quantitative HSV viral load data. B) All results with semiquantitative HSV viral load data. C) Correlation between quantitative and semiquantitative data for the 71 samples with numeric results (e.g., not “not detected” or “<250”) for both assays.

## DISCUSSION

After nearly 6 years as the only clinical genotypic acyclovir resistance assay available in the United States, we sequenced 136 samples providing important data on the landscape of acyclovir resistance in the US. We confirmed that most acyclovir resistance is due to frameshifts, particularly in homopolymer regions, that lead to early translation termination and production of non-functional UL23 (15). Of 93 unique mutations identified, we characterized 38 (40.9%) as causing acyclovir resistance (including the recently characterized G61E mutation), 32 (34.4%) as not associated with resistance, and identified 23 (24.7%) variants of uncertain significance (VUSs) without prior characterization. The identification of these VUSs highlights the need for additional biochemical characterization and continued sequencing to more fully sample the genetic diversity of HSV. Bluntly, there are many ways to break a protein and only through a combination of biochemical and sequencing efforts will we be able to more fully understand UL23 diversity and the effects of any clinically identified mutations. Despite over a decade of clinical HSV genotypic testing in Europe (6, 8, 10, 11), our identification of 23 VUSs—including 14 mutations neither previously characterized in the literature nor present in available GenBank UL23 sequences—highlights the potential for uncovering unique UL23 sequence variation in the US compared to Europe. Further comparisons of viral diversity between these two regions are warranted. Although biochemical characterization is necessary to fully understand the effects of the VUSs we identified on acyclovir resistance, sequencing context can provide insight. For instance, 7 previously uncharacterized mutations—Y4H, C6D, R41G, S74L, D108E, P223S, and E147G—were only present alongside known resistance mutations, making it difficult to predict their effects, but potentially indicating they were not strong resistance mutations on their own. Additionally, F368L, N301S, and the majority (4/5) of the A37S mutations were all found in a patient with multiple sequences who developed resistance through a frameshift mutation at site 147 about one month following their initial sequencing. Given the development of a clear resistance mechanism after initially identifying these substitutions, it is unlikely that they cause substantial acyclovir resistance. Conversely, P82L was present only in the context of mutations known not to be associated with acyclovir resistance (C6G, L42P, Q89R, A118V, V348I) and, as a mutation not previously seen, should be among the mutations prioritized for further characterization.

This study is limited by the relatively low number of sequenced samples (136), particularly compared to ordered tests (305), which highlights the challenges of running a genotypic resistance assay when there are no FDA-cleared HSV viral load assays and such testing is not always readily available to our outside clients to help triage samples for genotypic testing. Additionally, due to being a reference lab, clinical and detailed demographic information was not available for more than 80% of the samples tested. Our assay also only sequences *UL23* and not *UL30*. Although 95% of acyclovir- resistance mutations are due to UL23 mutations (15), not sequencing *UL30* means we may miss some resistant samples. As more antivirals are developed that target other HSV proteins (22, 23), additional sequencing targets and, potentially, whole genome sequencing may become necessary to fully characterize clinical HSV sequences as sensitive or resistant to available therapies (24–26). Such a transition could address another limitation of our assay in its use of Sanger sequencing rather than next-generation sequencing, which can more sensitively detect minor variants that could be associated with antiviral resistance (27, 28). In addition, with the potential availability of additional well-tolerated HSV antivirals, rapid identification of drug-resistant HSV becomes even more clinically actionable (29, 30). Although our turnaround time for genotypic testing of ∼10 days is substantially improved compared to phenotypic testing (7), further improvement in turnaround time will allow for more timely and, therefore, more impactful results. Nonetheless, genotypic testing will always rely on accurate variant interpretation, and these clinical efforts will necessarily happen hand-in-hand with continued biochemical and phenotypic characterization of HSV mutations to ensure we can provide accurate clinical interpretations for identified variants.

## Acknowledgements

ALG reports contract testing to UW from Abbott, Cepheid, Novavax, Pfizer, Janssen, Assembly Biosciences, Aicuris, Innovative Molecules, and Hologic, research support from Gilead, personal fees from Arisan Therapeutics, outside of the described work. The remaining authors report no conflicts of interest.

